# Differential processing and habituation in distinct spatial frequency channels in V1 of a mouse model of fragile X syndrome

**DOI:** 10.1101/2020.01.24.919035

**Authors:** Alexandr Pak, Samuel T. Kissinger, Alexander A. Chubykin

## Abstract

Extraction of both common and unique features across different visual inputs is crucial for animal survival. Regularities in the visual input lead to learning of the general principles governing an environment, whereas unique features are important for novelty detection. Low and high spatial frequencies (SF) represent two different channels of visual perception, which may be playing different roles in the processing of global pattern and local details. Alterations in the processing of these different SF channels may lead to impaired visual perception. Excessive detail-oriented processing and reduced habituation to sensory stimuli are some of the hallmarks of altered sensory perception in autism. However, the underlying neural mechanisms of these impairments are not understood. To gain insight into the pathophysiology of these impairments, we investigated the low and high SF channels in V1 of Fmr1 KO mice, the mouse model of Fragile X syndrome (FX). We first provide behavioral evidence for reduced habituation of both pupillary baseline and surprise responses in FX mice. Using silicon probe recordings, we demonstrate excessive processing of high SF stimuli in the late stages of visual responses in V1 of FX mice. We then show a reduced adaptation during a visual oddball paradigm in neurons preferring low but not high SF. Overall, our findings suggest that altered processing in distinct SF channels might contribute to altered visual perception and learning in FX and autism.

## Introduction

Fragile X Syndrome (FX) is the most common cause of intellectual disability and of the inherited form of autism. Nearly 1 in 4000 males and half as many females are affected by this condition. It is associated with social communication deficits, hyperactivity, and sensory hypersensitivity (Freund and Reiss, 1991). Given the comorbidity of FX and autism, *Fmr1* KO mice (FX mice) represent a well-defined genetic model that can provide neural circuit-level insights into autism, especially considering the vast diversity of phenotypes and manifestations observed in autism spectrum disorders (ASDs). Such diverse alterations posit a challenge to develop effective diagnostic and treatment tools. FX mice have been shown to exhibit cellular, circuit, and behavioral alterations that recapitulate some of the manifestations observed in human individuals with FX. Prior autism research has been mostly focused on social-cognitive and behavioral impairments (Robertson and Baron-Cohen, 2017). However, a recent revision of diagnostic criteria for autism recognized sensory processing as an important factor to be considered (American Psychiatric Association, 2013). Previous research in humans suggests that sensory alterations may be predictive of social communication deficits later in life in autism (Boyd et al., 2010; Turner-Brown et al., 2012).

Both human and animal studies provide evidence that there is impaired information processing in early sensory areas in both autism and FX (Goel et al., 2018; Rais et al., 2018). Sensory hypersensitivity and reduced adaptation to sensory stimuli are some of the hallmark perceptual impairments in autism. An increase in visual detail processing is often reported in this condition. Visual oddball paradigm studies revealed reduced habituation to repeated stimuli and novel distractors in autistic patients (Sokhadze et al., 2017). Similarly, alterations in the event related potentials (ERPs) during the auditory and visual oddball tasks were found in FX patients (Van der Molen et al., 2012). Recent work in FX mice found circuit-level impairments in early visual processing, including reduced orientation tuning and functional output from fast-spiking neurons in V1. Reduced orientation tuning of the neurons in the visual cortex correlated with the decreased ability to resolve different orientations of sinusoidal grating stimuli in both mice and human individuals with FX (Goel et al., 2018). Furthermore, altered dendritic spine function and integration were found in early sensory areas in FX mice (Domanski et al., 2019). Structural and functional imaging studies of FX mice revealed local hyperconnectivity and long-range hypoconnectivity in V1. These studies suggest that primary cortical areas may be functionally decoupled from other brain regions, giving rise to the impaired high-level modulation of sensory processing (Haberl et al., 2015). Overall, these studies suggest that there may be circuit-level impairments in early sensory processing in FX.

To shed light on the neural basis of atypical visual perception in FX, we investigated how statistical context influences visual information processing by testing both basic and contextual processing of spatial frequencies (SF) in V1 of FX mice. We measured visually evoked potentials (VEPs) and unit responses in a SF oddball paradigm (Hamm and Yuste, 2016; Ulanovsky et al., 2003). Two stimuli were presented at different probabilities so that one was a standard stimulus (STD, frequent, redundant), which builds a statistical context. Another one was rare and violated the expectations of the STD stimulus leading to a mismatch response. This response is believed to reflect a perceptual deviance or change detection. First observed in EEG studies in humans as delayed negative deflection in event-related potentials, later called mismatch negativity (MMN) (Naatanen et al., 1978), it has been replicated in different species and sensory modalities (Chen et al., 2015; Musall et al., 2015; Parras et al., 2017). A decrease in the neural response to the standard stimulus (STD) may be attributed to the predictability of the stimulus because the incoming sensory input matches prediction. Alternatively, it may also be explained by the presynaptic short-term plasticity mechanisms. On the other hand, the upregulation of deviant (DEV) responses is due to the violation of the expectations. It triggers a sensory prediction error (PE) signal, which is defined as the difference between sensory input and its prediction (Stefanics et al., 2014). Stimulus specific adaptation (SSA) was defined as the difference between CTR and STD, whereas PE was the difference between DEV and CTR stimuli (Hamm and Yuste, 2016; Parras et al., 2017).

Here, we performed silicon probe recordings in WT and FX mouse V1 during the SF tuning and the oddball paradigm. We investigated the basic sensory processing of SF and the contextual modulation of neurons preferring various SF bands. Firstly, we demonstrate reduced habituation during visual oddball paradigm as indicated by baseline pupil and surprise response in FX mice. Second, we report excessive processing in late neural responses, especially at high SF. Third, we show a reduced adaptation and enhanced prediction errors in low SF channels. Overall, our findings suggest that differential alterations in low and high spatial frequency channels may contribute to altered sensory perception in FX and autism.

## Results

### Pupillometry reveals reduced habituation and surprise optimization in FX animals

Pupil dilation in response to surprising events may represent a physiological proxy for prediction error and acute changes in the arousal state of an animal (Preuschoff et al., 2011). Given that reduced habituation and adaptation are reported in human individuals with FX, we first tested whether FX mice exhibit similar impairments in visually evoked behaviors by recording pupillometry during a 2-session oddball paradigm (**Figure 1A and B**). We used deviant stimuli that were drastically different from standard stimuli to induce a strong surprise response. A checkerboard stimulus was used as a deviant pattern and sinusoidal grating as a standard. We observed a strong surprise response to DEV stimuli in both groups (**Figure 1C**, session 1, STD vs DEV: WT (P = 1.61E-7), n = 18 mice, FX (P = 1.63E-7), n = 21 mice, Mann-Whitney U test). We ran the oddball sequence for two sessions with at least a 60 s break in between. We then analyzed how the baseline pupil size and surprise response were different in session 1 vs 2 in WT and FX mice. In session 2, we saw a significant decrease in both baseline pupil size and the surprise response in WT mice (**Figure 1D**, WT session 1 vs 2: baseline (0-1s) (P = 0.003), n = 13 mice; surprise (1-4 s) (P = 0.0003), n = 13 mice, Paired t-test). Interestingly, we did not observe a significant decrease in either baseline pupil size or the surprise response in FX mice (**Figure 1D** right, FX session 1 vs 2: baseline (0-1s) (P = 0.76), n =17 mice; surprise (1-4 s) (P = 0.33), n = 17 mice, Paired t-test). These results suggest that WT, unlike FX mice, demonstrate habituation of visually evoked pupil increases after an experience with the oddball paradigm.

**Figure 1.**
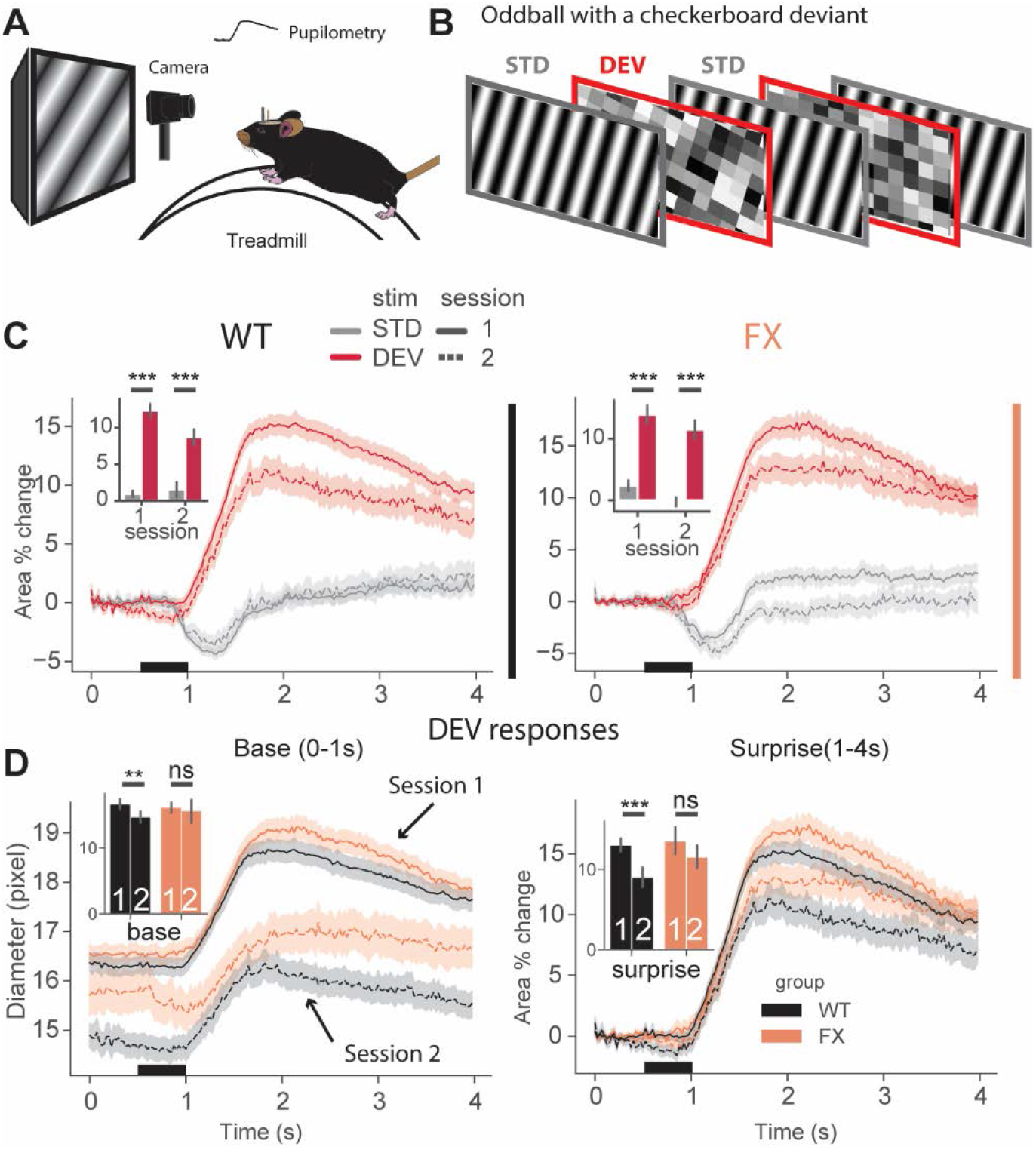
Pupillometry during a visual oddball paradigm reveals reduced habituation and surprise adaptation in FX mice. **A.** Experimental setup for pupillometry recordings in awake head-fixed mice. **B.** Oddball paradigm where gratings serve as standard (STD) visual stimulus while checkerboards serve as deviant stimuli (DEV) to induce strong surprise responses. **C.** Both wild type (WT) and fragile X (FX) mice show strong surprise responses to DEV (red) stimuli. **D.** Left panel: baseline pupil diameter during deviant trials in WT and FX mice. WT but not FX mice habituate to this paradigm after the first recording session as indicated by decrease in baseline pupil size. Right panel: pupillary surprise responses during deviant trials in WT and FX mice. Surprise responses decrease in the 2^nd^ session in WT but not in FX mice. Numbers inside the bar plot represent the session number. Error bars indicate s.e.m. \**p* < 0.05, ** *p* < 0.01, \*\*\**p* < 0.001.

### Excessive processing of high spatial frequencies in V1 of FX mice in late unit responses

We then investigated the neural correlates underlying reduced habituation in FX, we performed *in vivo* extracellular electrophysiology in awake head-fixed mice (**Figure 2A, B**). Using 64 channel silicon probes that span the cortical depth of V1 (Shobe et al., 2015b), we investigated the laminar specific visual processing of different spatial frequencies (SF) (**Figure 3A**). We presented animals with SF filtered visual noise stimuli using six different non-overlapping SF bands to probe both low and high SF processing streams (**Figure 2C,D**). We first analyzed the SF tuning of single units across the cortical column, revealing a differential strength and time course of the responses in the population firing rate (**Figure 3A**). Tuning curves for different layers were then constructed by averaging the baseline-corrected firing rate within 0.05-0.2s relative to the stimulus onset of all units within a particular layer. We found upregulated responses in superficial layers of FX mice (**Figure 3B**, superficial WT vs FX: SF: 7.5E-3 cpd (P = 0.04), SF: 0.015 cpd (P = 0.004), SF: 0.03 cpd (P = 0.04), SF: 0.06 cpd (P = 0.009), SF: 0.12 cpd (P = 0.005), and SF: 0.24 cpd (P = 0.009), Mann-Whitney U test, p-values were adjusted for multiple comparisons using Benjamini-Hochberg method, n = 357 vs 233 units), while no differences could be found between WT and FX in layer 4 or deep layers (**Figure 3B**). The time course of SF tuning revealed enhanced activity in late neural responses in all layers, especially at higher SF (**Figure 3C**). To investigate the time course of SF processing, we averaged firing rates within different time windows. We found a significantly stronger response at higher SF (>0.06 cpd) in late intervals, 0.25 s after stimulus onset (**Figure 3D**, mean firing rate 0.25-0.35 s WT vs FX: SF: 7.5E-3 cpd (P = 0.15), SF: 0.015 cpd (P = 0.03), SF: 0.03 cpd (P = 0.15), SF: 0.06 cpd (P = 0.02), SF: 0.12 cpd (P = 0.002), and SF: 0.24 cpd (P = 0.01); mean firing rate 0.35-0.45 s WT vs FX: SF: 7.5E-3 cpd (P = 0.15), SF: 0.015 cpd (P = 0.01), SF: 0.03 cpd (P = 0.28), SF: 0.06 cpd (P = 0.15), SF: 0.12 cpd (P = 0.006), and SF: 0.24 cpd (P = 0.008), Mann-Whitney U test adjusted for multiple comparisons, 1057 vs 820 units). Next, we performed SF neural decoding using population spike counts (**Figure 3E**). We trained classifiers on spike counts from different time windows of WT and FX mice using a linear discriminant analysis with a 4-fold cross-validation with 5 repeats. Classifiers trained on spike counts from 0.05-0.5 s performed similarly (SF classification error WT vs FX: 9% vs 12%). WT classifiers performed slightly better in early time windows (SF classification error WT vs FX 0.05-0.15 s: 16% vs 23%; 0.15-0.25 s: 6% vs 10%). However, classifiers trained on the intervals after 0.25 s show a reduced error in FX vs WT mice (SF classification error WT vs FX 0.25-0.35 s: 22% vs 16%; 0.35-0.45 s: 26% vs 15%), suggesting enhanced processing in late neural responses. Together, these findings suggest an enhancement of processing in late neural responses in FX vs WT mice, especially at high spatial frequencies.

**Figure 2.**
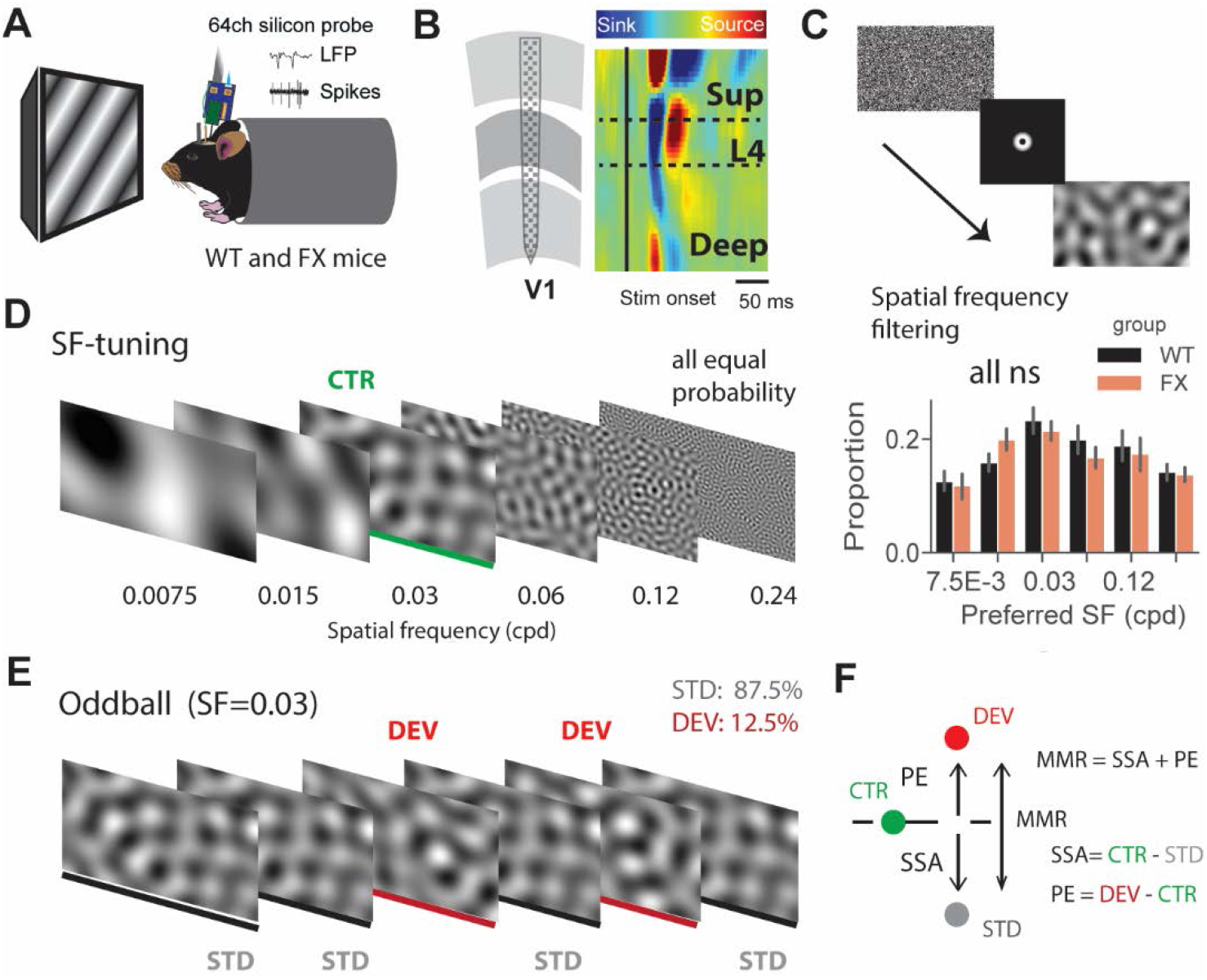
A visual oddball paradigm with all the stimuli containing the same low level features (spatial frequency) but different global spatial frequency patterns and expectancy. **A.** In vivo extracellular silicon probe recordings in V1 of head-fixed mice. **B.** Schematic of a 64 channel silicon probe spanning the whole cortical depth and an example current source density (CSD) heat map. **C.** To generate visual stimuli, we performed spatial frequency (SF) filtering of white noise. **D.** Left: we used 6 different non-overlapping SF bands from 7.5E-3 to 0.24 cpd for spatial frequency tuning. Stimuli were presented in a pseudorandom order and had equal probability. Right: proportion of neurons preferring various SF in WT and FX mice. **E.** The oddball sequence contained stimuli of the same SF (0.03 cpd) that only differ in their probability and overall texture. Standard (STD) and deviant (DEV) stimuli were presented with a probability of .875 and 0.125, respectively. Note: Since both STD and DEV had the same SF, neurons preferring the target SF are expected to show a stronger adaptation compared to those not preferring it. **F.** We decomposed a neuronal mismatch response into stimulus-specific adaptation (SSA) and prediction error (PE) by comparing STD and DEV responses to the control (CTR) from the SF tuning sequence. CTR served as the baseline response for SF of 0.03 cpd. Error bars indicate s.e.m. *\*p* < 0.05, \*\**p* < 0.01, \*\*\**p* < 0.001.

**Figure 3.**
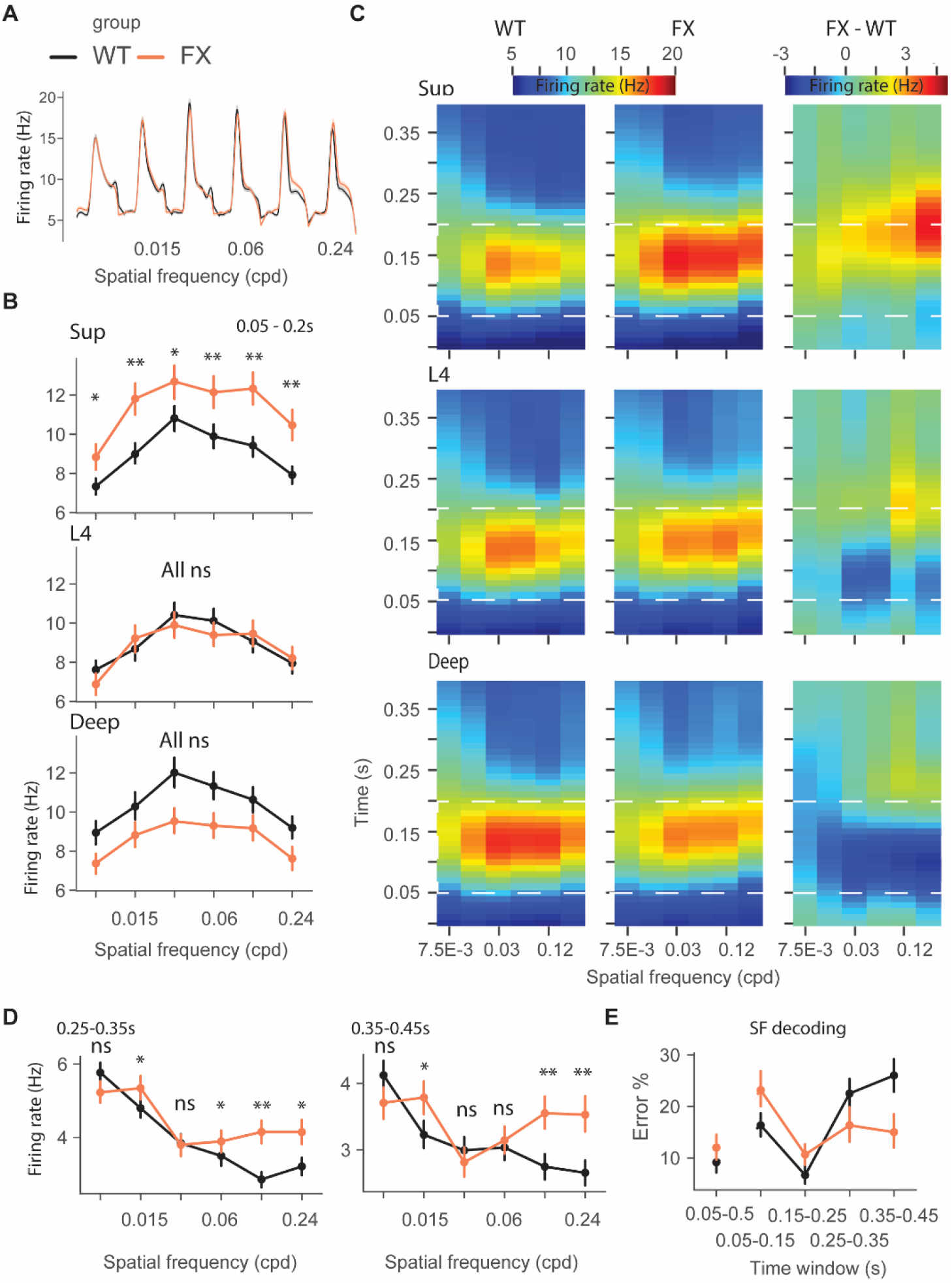
Excessive processing of high SF stimuli in late responses of single units in FX mice. **A.** Population average firing rates of all units in response to the SF tuning sequence. **B.** SF tuning across different cortical layers. The mean firing rate within 0.05-0.2 s relative to stimulus onset of visual stimulation was computed, and neural responses were averaged across units to create the point plots **C.** Time course analysis of SF tuning across the cortical layers. Unit responses for different SF stimuli were plotted for each time step to create the heatmaps. **D.** Population mean SF tuning responses were averaged for different time intervals. Averaging responses within 0.25-0.35 s and 0.3-0.45 s revealed significantly stronger firing rates at higher spatial frequencies in FX animals. **E**. Population spike counts from different time windows were used for SF neural decoding. The classifier was trained on responses after 0.25 s relative to stimulus onset had lower error in FX vs WT mice. Error bars indicate s.e.m. \**p* < 0.05, \*\**p* < 0.01, \*\*\**p* < 0.001.

### Altered oscillatory dynamics during a visual oddball paradigm in FX mice

To efficiently adapt to the environment, unexpected events should give rise to specific prediction errors that would result in the adjustment of expectations. To study this process, we sought to design a visual oddball paradigm such that the low-level features (spatial frequency) of visual stimuli were the same, but only the contextual information including the stimulus texture or probability would modulate responses. We used filtered noise stimuli where a commonlypresented standard (STD) stimulus differs in texture from a sparsely presented deviant (DEV) stimulus (**Figure 2C,E, F**). Both STD and DEV stimuli had the same SF of 0.03 cpd. To control for both SF and probability, responses to the same SF of 0.03 cpd from the SF tuning experiment were used as controls (CTR). This allowed us to investigate how common and unique features are processed in different SF channels in WT and FX mice. Since STD and DEV stimuli were of the same SF, we expected a strong adaptation in neurons preferring SF of 0.03 cpd.

We first investigated local field potential (LFP) responses during the oddball sequence, which represent local population subthreshold activity. Layer 4 LFP mean population traces in response to STD, DEV, and CTR are shown in **Figure 4A**. The strongest responses were selected from channels within the window of 0.05-0.2 s relative to the stimulus onset. Responses were then averaged for each recording. DEV and CTR responses were stronger than STD, likely due to visual adaptation to the STD driven by the higher probability of presentation. We initially expected DEV stimuli to evoke stronger responses than CTR as was previously found (Hamm and Yuste, 2016; Parras et al., 2017). However, no significant differences were observed between DEV and CTR probably due to the SF adaptation given that DEV shared SF with STD (**Figure 4A, B**, L4 VEP amplitude WT: STD vs DEV (P = 0.02), STD vs CTR (P = 0.007), and DEV vs CTR (P = 0.98), n = 24 recordings; FX: STD vs DEV (P = 0.02), STD vs CTR (P = 0.01), DEV vs CTR (P = 0.98), n = 21 recordings, Mann-Whitney U test adjusted for multiple comparisons). In later parts of the study, we show that the magnitude of adaptation and prediction errors depends on the preferred SF of the neurons. Direct comparisons of stimuli between groups did not reveal any differences. We next performed a time-frequency analysis using a complex wavelet convolution (**Figure 4C**). Power was normalized to the baseline and averaged within 0.05-0.2 s across different frequencies. We found a significant theta modulation in FX but not in WT animals (**Figure 4D**, theta WT: STD vs DEV (P = 0.15), STD vs CTR (P = 0.15), and DEV vs CTR (P = 0.32), n = 24; FX: STD vs DEV (P = 0.04), STD vs CTR (P = 0.04), and DEV vs CTR (P = 0.2), n = 21 recordings, Mann-Whitney U test adjusted for multiple comparisons). Alpha oscillations were significantly decreased in response to STD in both groups (**Figure 4B**, alpha WT: STD vs DEV (P = 0.04), STD vs CTR (P = 0.02), and DEV vs CTR (P = 0.35), n = 24; FX: STD vs DEV (P = 0.09), STD vs CTR (P = 0.03), and DEV vs CTR (P = 0.16), n = 21 recordings, Mann-Whitney U test adjusted for multiple comparisons) and were significantly stronger for DEV in WT mice. There were no differences in beta oscillations. Low gamma oscillations were significantly modulated in WT mice (**Figure 4B**, low gamma WT: STD vs DEV (P = 0.05), STD vs CTR (P = 0.002), and DEV vs CTR (P = 0.16), n = 24; FX: STD vs DEV (P = 0.09), STD vs CTR (P = 0.05), and DEV vs CTR (P = 0.34), n = 21 recordings, Mann-Whitney U test adjusted for multiple comparisons). Decreased high gamma in response to STD was observed in both groups. However, it was only significantly upregulated in response to DEV in WT mice (**Figure 4B**, high gamma WT: STD vs DEV (P = 0.0021), STD vs CTR (P = 0.006), and DEV vs CTR (P = 0.36), n = 24; FX: STD vs DEV (P = 0.14), STD vs CTR (P = 0.006), and DEV vs CTR (P = 0.14), n = 21 recordings, Mann-Whitney U test adjusted for multiple comparisons). A direct comparison of stimuli between groups did not reveal any differences across different frequency bands. Overall, we found altered oscillatory activity in response to DEV stimulus in FX mice.

**Figure 4.**
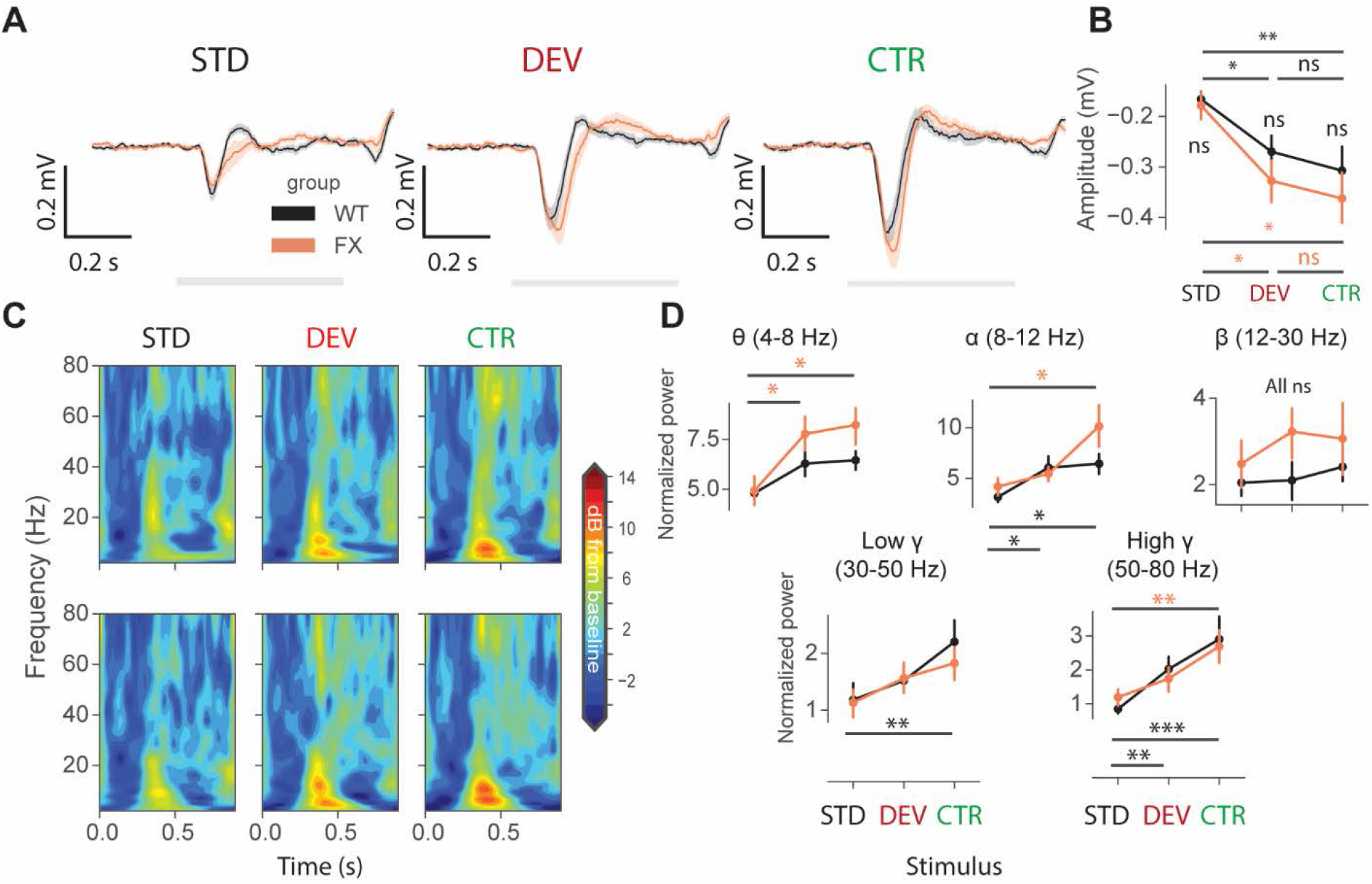
Altered oscillatory dynamics in FX mice during a visual oddball paradigm. **A.** Averaged layer 4 LFP traces in response to STD, DEV, and CTR stimuli. **B.** The point plot shows the mean and s.e.m. of the strongest negative amplitude within 0.05-0.2 s relative to the stimulus onset. **C.** Time frequency spectra of the L4 LFP traces of WT (top) and FX (bottom). **D.** Point plots show the mean power within 0.05-0.2 s relative to the stimulus onset across different frequency bands. Error bars indicate s.e.m. \**p* < 0.05, \*\**p* < 0.01, \*\*\**p* < 0.001.

### Reduced adaptation and enhanced prediction errors in FX mice following the visual oddball

Next, we focused on single-unit activity during the oddball paradigm by investigating how neurons preferring different SF were modulated by SSA and PEs. We first identified the preferred SF for each neuron as the one that induces a maximum response, mean firing rate between 0.05-0.2 s. We only selected units that were excited by visual stimuli. We first investigated oddball responses in the low spatial frequency channels (**Figure 5A and B**). Units preferring the lowest tested SF of 7.5E-3 cpd showed a qualitative difference between groups and stimuli but did not reach significance after correcting for multiple comparisons (**Figure 5A**, all comparisons p > 0.05, n = 65 and 46 units). Note that both STD and DEV were stronger in FX mice. Neurons preferring SF of 0.015 cpd showed a significant adaptation in WT but not in FX mice. DEV was significantly stronger than CTR only in FX but not in WT animals (**Figure 5B**, WT: STD vs DEV (P = 0.0002), STD vs CTR (P = 0.0052), DEV vs CTR (0.21), n = 108 units; FX: STD vs DEV (P = 0.0001), STD vs CTR (P = 0.1), and DEV vs CTR (P = 0.0085), n = 111 units, Mann-Whitney U test adjusted for multiple comparisons). A direct comparison between groups revealed significantly stronger responses to STD and DEV in FX mice (WT vs FX: STD (P = 0.0019), DEV (P = 0.0009), and CTR (P = 0.10), n = 108 vs 111 units, Mann-Whitney U test adjusted for multiple comparisons).

**Figure 5.**
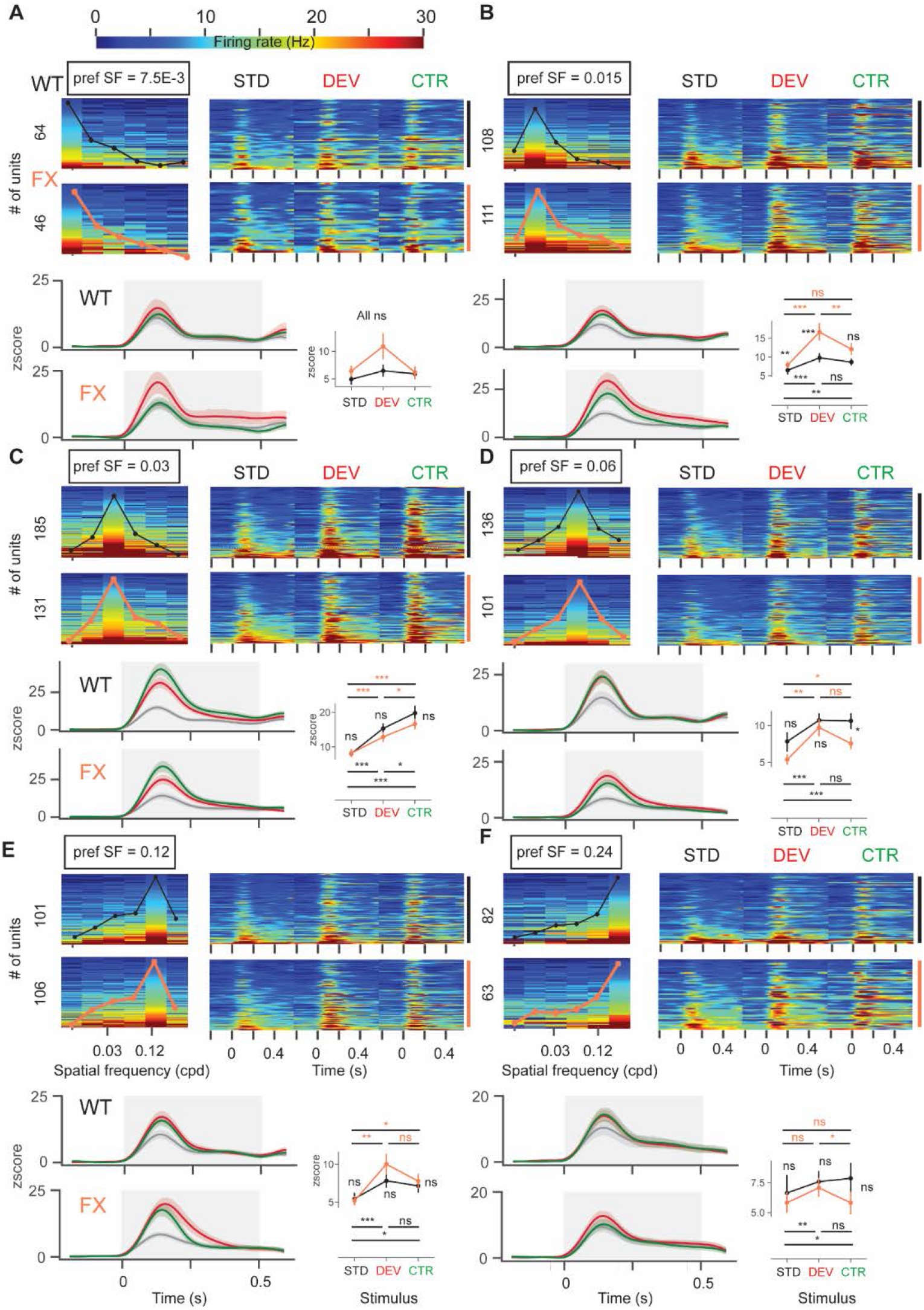
Reduced adaptation and enhanced prediction errors in neurons preferring low SF in FX mice compared to WT. **A.** Neural responses to the oddball paradigm of low spatial frequency preferring neurons. Units were grouped by their preferred SF, which was defined as a maximum response in the tuning curve. Left: preferred SF=0.0075: a heatmap of single-unit tuning curves is overlaid with a population mean curve. Three additional heatmaps show single-unit firing rates in response to STD, DEV, and CTR stimuli. The line plots below represent mean z-scored responses of the units from the heatmap. Point plots show the mean ± s.e.m. of the z-scored firing rate between 0.05-0.5 s relative to the stimulus onset. Right: preferred SF = 0.015 cpd. **B.** Same as in A, but preferred SF = 0.015 cpd. **C.** Same as in A, but preferred SF = 0.03 cpd. **D.** Same as in A, but preferred SF = 0.06 cpd. **E.** Same as in A, but preferred SF = 0.12 cpd. **F.** Same as in A, but preferred SF = 0.24 cpd. Error bars indicate s.e.m. \**p* < 0.05, \*\**p* < 0.01, \*\*\**p* < 0.001.

An identical analysis was performed on oddball responses to the intermediate spatial frequencies of 0.03 and 0.06 cpd (**Figure 5C and D**). Units that preferred the target oddball SF of 0.03 cpd showed significant adaptation in both groups. Furthermore, CTR was significantly stronger than DEV in this neural subpopulation in both genotypes (**Figure 5C**, WT: STD vs DEV (P < 0.0001), STD vs CTR (P < 0.0001), DEV vs CTR (P = 0.01), n = 186 units; FX: STD vs DEV (P = 0.00002), STD vs CTR (P < 0.0001), and DEV vs CTR (P = 0.01), n = 132 units, Mann-Whitney U test adjusted for multiple comparisons). This was expected due to the STD and DEV having the same SF. A direct comparison of stimuli between groups did not reveal any difference. For neurons preferring SF of 0.06 cpd, both groups showed adaptation. DEV was not different from CTR (**Figure 5D**, WT: STD vs DEV (P = 0.00003), STD vs CTR (P = 0.0006), DEV vs CTR (P = 0.43), n = 140 units; FX: STD vs DEV (P = 0.00025), STD vs CTR (P = 0.01), and DEV vs CTR (P = 0.19), n = 101 units, Mann-Whitney U test adjusted for multiple comparisons). A direct comparison between groups revealed a significantly weaker CTR response in FX mice (**Figure 5D**, WT vs FX: STD (P = 0.22), DEV (P = 0.22), and CTR (P = 0.04), n = 140 vs 101 units, Mann-Whitney U test adjusted for multiple comparisons).

We then focused on neurons preferring high spatial frequencies (**Figure 5E and F**). Both groups showed adaptation in neurons that prefer SF of 0.12 cpd. DEV was not different from CTR responses (**Figure 5E**, WT: STD vs DEV (P = 0.0009), STD vs CTR (P = 0.01), DEV vs CTR (P = 0.19), n = 112 units; FX: STD vs DEV (P = 0.001), STD vs CTR (P = 0.02), and DEV vs CTR (P = 0.19), n = 106 units, Mann-Whitney U test adjusted for multiple comparisons). A direct comparison did not reveal any differences between groups. Units that prefer SF of 0.24 cpd showed adaptation only in WT animals. DEV was significantly stronger than CTR only in FX mice (**Figure 5F**, WT: STD vs DEV (P = 0.009), STD vs CTR (P = 0.04), DEV vs CTR (P = 0.22), n = 84 units; FX: STD vs DEV (P = 0.07), STD vs CTR (P = 0.31), and DEV vs CTR (P = 0.04), n = 63 units, Mann-Whitney U test adjusted for multiple comparisons). A direct comparison of stimuli between groups did not reveal any differences. Overall, we observed a strong adaptation and significantly weaker DEV vs CTR responses in neurons preferring SF of 0.03 cpd. This result was expected given that STD and DEV shared the same SF. Reduced adaptation and significant PEs were only found in neurons preferring SF of 0.015 and 0.24 cpd. Significant differences in STD and DEV responses between groups were only found in units preferring SF of 0.015 cpd.

### Reduced population stimulus-specific adaptation and increased prediction errors in the low SF channel in FX mice

We next investigated how neurons grouped by their preferred SF are tuned to the STD, DEV, and CTR stimuli. We averaged the baseline subtracted firing rate across different layers and found that L4 of both groups showed feature-specific responses to all 3 stimuli, with tuning curves exhibiting similar shapes and peaks. Interestingly, superficial and deep layers of FX mice did not exhibit SF group-specific tuning of both STD and DEV responses (**Figure 6A**). Then, we focused on the population mean iSSA and iPE indices across neuronal groups preferring different spatial frequencies:

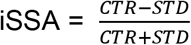

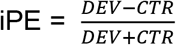

**Figure 6.**
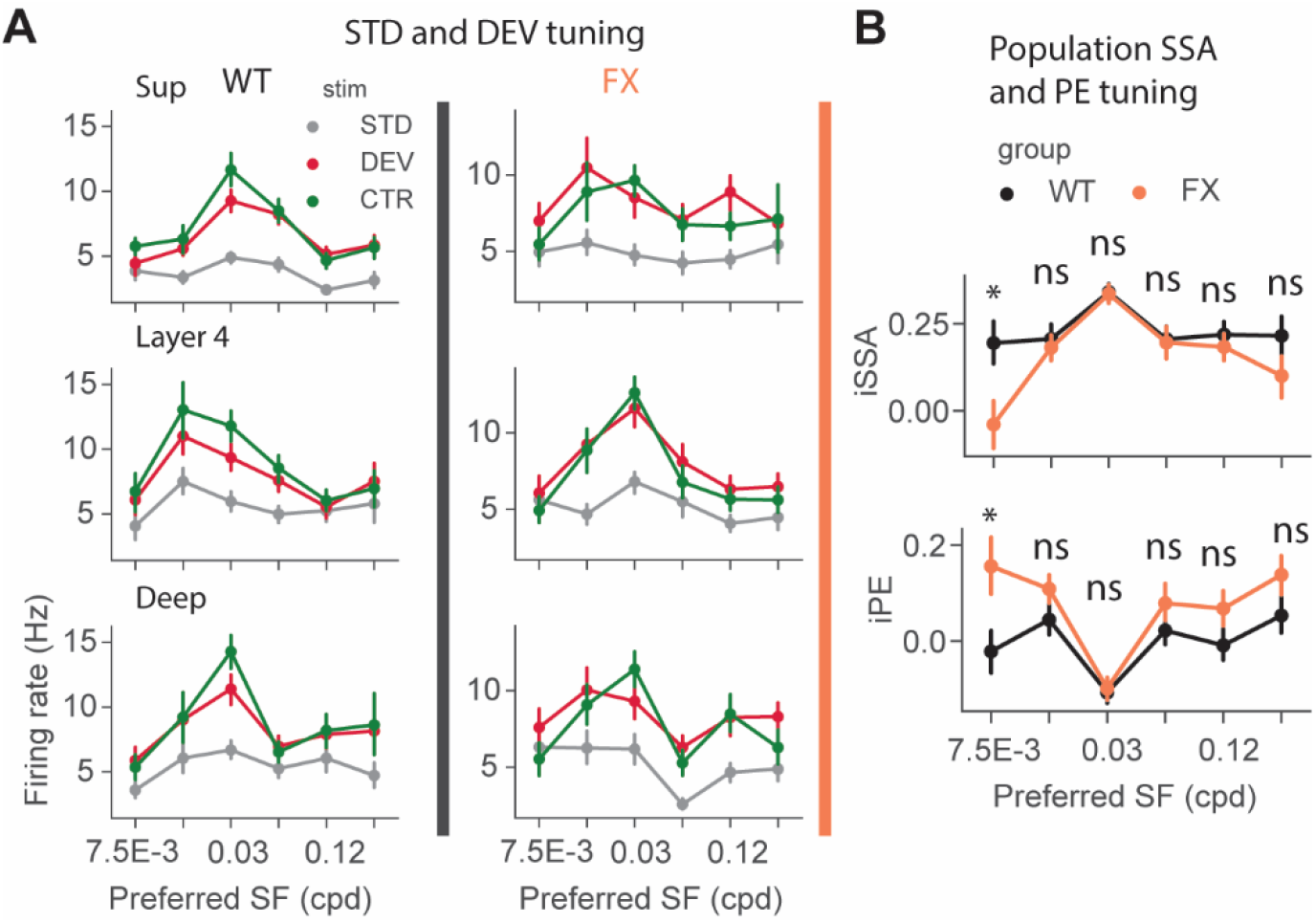
Adaptation and prediction error tuning across neural populations preferring different spatial frequencies. **A.** Neural responses to STD, DEV, and CTR of units preferring various SF in WT (left) and FX animals (right). Note: Feature specificity of STD and DEV stimuli in WT but not in FX mice in superficial and deep layers. **B.** Mean of iSSA and iPE across different neural subpopulations preferring different SF in WT vs FX mice. iSSA tuning peaks at the target SF (0.03 cpd) in both genotypes. However, the iSSA is significantly lower at the SF of 7.5E-3 cpd in FX vs WT mice. iPE curves have a similar shape in both groups, with a negative peak at target oddball SF of 0.03 cpd. iPE is significantly stronger at the SF of 7.5E-3cpd in FX mice. Error bars indicate s.e.m. \**p* < 0.05, \*\**p* < 0.01, \*\*\**p* < 0.001.

We found differences at the left tail of the adaptation tuning curve in WT vs FX animals. In particular, we found a significant difference in units preferring the lowest SF tested (**Figure 6B** top, iSSA WT vs FX: SF: 7.5E-3 cpd (P = 0.029), n = (51, 37 units); SF: 0.015 cpd (P = 0.46), n = (88, 107); SF: 0.03 cpd (P = 0.47), n = (168, 120 units); SF: 0.06 cpd (P = 0.47), n = (121, 84 units); SF: 0.12 cpd (P = 0.46), n = (92, 88 units); SF: 0.24 cpd (P = 0.22), n = (67, 50 units), Mann-Whitney U test, p-values were adjusted for multiple comparisons using Benjamini-Hochberg method). Feature tuning of prediction errors was also altered in FX animals. PEs had a negative peak at target oddball SF=0.03 cpd in both groups, consistent with strongest SSA in those neurons. Interestingly, PEs were upregulated in units preferring low but not high SF (**Figure 6B** bot, iPE WT vs FX: SF: 7.5E-3 cpd (P = 0.01), SF: 0.015 cpd (P = 0.10), SF: 0.03 cpd (P = 0.26), SF: 0.14 cpd (P = 0.14), SF: 0.12 cpd (P = 0.10), SF: 0.24 cpd (P = 0.10), same number of units for each comparison as for iSSA, Mann-Whitney U test adjusted for multiple comparisons). Together, these results suggest that there is a reduced adaptation and enhanced prediction errors in the low SF channel of FX mice.

### Prediction errors are present across the cortical column in FX but not in WT mice

We found the strongest adaptation in neurons preferring the target oddball SF of 0.03 cpd, which is aligned with our predictions. To isolate true population prediction errors across the cortical column, we focused on units that did not prefer SF 0.03. We investigated the SSA and PE across different cortical layers. **Figure 7A and B** show the example unit responses to STD, DEV, and CTR stimuli from WT and FX mice. In superficial layers, we found that WT but not FX mice show significantly stronger responses to CTR vs STD. On the other hand, only FX mice show significantly stronger response to DEV vs CTR (**Figure 7E** top, WT: STD vs DEV (P < 0.0001), STD vs CTR (P < 0.0001), DEV vs CTR (P = 0.21), n = 219 units; FX: STD vs DEV (P = 0.0001), STD vs CTR (P = 0.056), and DEV vs CTR (P = 0.03), n = 152 units, Mann-Whitney U test, p-values were adjusted for multiple comparisons using Benjamini-Hochberg method). A direct comparison between groups reveal significantly stronger responses to STD and DEV in FX vs WT mice (**Figure 7E** top, WT vs FX: STD (P = 0.00038), DEV (P = 0.03), and CTR (P = 0.35), n = 219 vs 152 units). Furthermore, iSSA but not iPE distributions were significantly different in FX vs WT mice (**Figure 7F** top, WT vs FX: iSSA (P = 0.02); iPE (P = 0.055), n = 181 vs 134 units, Kolmogorov-Smirnov 2 sample test).

**Figure 7.**
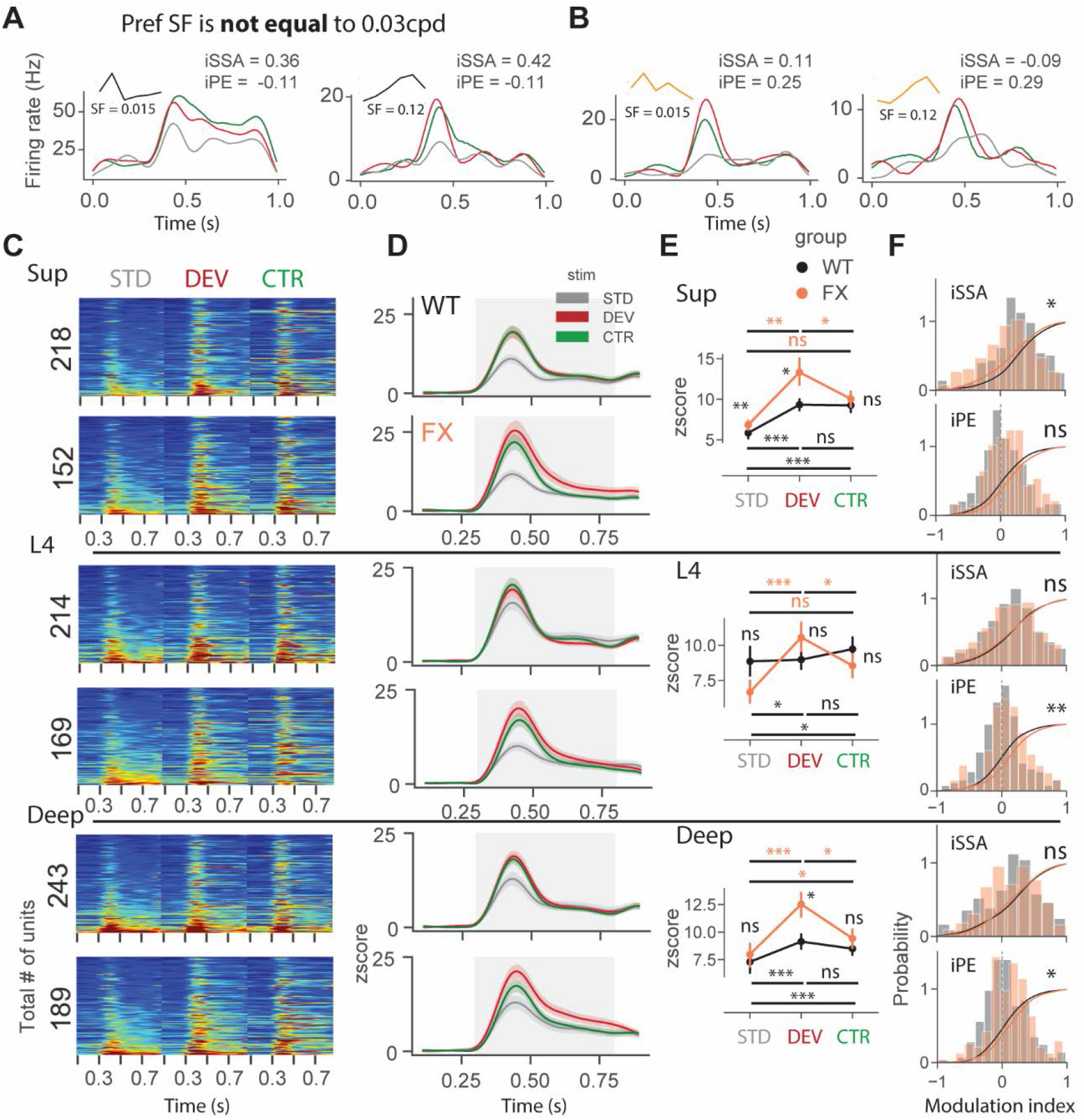
Prediction errors are present across the whole cortical column in FX but not WT animals. **A.** Example single unit responses to STD, DEV, and CTR stimuli from WT mice. Inset: the SF tuning curves for each unit and their preferred SF. **B.** Same as in A but from FX mice. **C.** Heatmaps of single unit firing rates in response to STD, DEV, and CTR stimuli across different layers from WT (top) and FX (bottom) mice. **D.** Line plots showing the mean response of the units in the heatmaps. **E.** Point plots show the mean and s.e.m. of the responses between 0.35-0.8 s of the three different stimuli. **F.** Cumulative distribution functions are overlaid with histograms of iSSA (top) and iPE (bottom) distributions across different cortical layers. Error bars indicate s.e.m. \**p* < 0.05, \*\**p* < 0.01, \*\*\**p* < 0.001.

Unit responses from L4 showed a similar trend. There was a significant adaptation in WT but not in FX mice. DEV responses were significantly stronger only in FX mice **Figure 7E** middle, WT: STD vs DEV (P = 0.02), STD vs CTR (P = 0.02), DEV vs CTR (P = 0.44), n = 215 units; FX: STD vs DEV (P = 0.0004), STD vs CTR (P = 0.09), and DEV vs CTR (P = 0.04), n = 169 units, Mann-Whitney U test, p-values were adjusted for multiple comparisons using Benjamini-Hochberg method). A direct comparison of stimuli between groups did not reveal any differences. The distributions of iPE but not iSSA indices were significantly different between groups (**Figure 7F** middle, WT vs FX: iSSA (P = 0.91); iPE (P = 0.003), n = 182 vs 138 units, Kolmogorov-Smirnov 2 sample test).

An analysis of deep layer responses revealed a significant adaptation in both groups. However, similarly to superficial layers and L4, DEV was significantly stronger than CTR only in FX mice (**Figure 7E** bottom, WT: STD vs DEV (P < 0.0001), STD vs CTR (P = 0.0006), DEV vs CTR (P = 0.14), n = 243 units; FX: STD vs DEV (P < 0.0001), STD vs CTR (P = 0.01), and DEV vs CTR (P = 0.01), n = 190 units, Mann-Whitney U test, p-values were adjusted for multiple comparisons using Benjamini-Hochberg method). Furthermore, a direct comparison of stimuli between groups revealed a stronger DEV response in FX mice (**Figure 7E** bottom, WT vs FX: STD (P = 0.06), DEV (P = 0.01), and CTR (P = 0.15), n = 243 vs 190 units). iPE but not iSSA distributions were significantly different between groups in deep layers (**Figure 7F** bottom, WT vs FX: iSSA (P = 0.41); iPE (P = 0.03), n = 204 vs 171 units, Kolmogorov-Smirnov 2 sample test). Together, our findings suggest that there are a reduced adaptation and increased prediction errors in neurons not preferring target oddball SF across the laminar depth in FX mice.

## Discussion

The lack of a common framework to explain the disparate sensory and social-cognitive deficits in FX and autism is a major roadblock to scientific progress and designing effective diagnostic and intervention tools. Atypical sensory processing has recently been recognized to be an important diagnostic criterion for autism (American Psychiatric Association, 2013). Furthermore, early sensory alterations are predictive of social communication deficits later in life (Robertson and Baron-Cohen, 2017). Investigating the reproducible sensory perception paradigms in well-defined genetic models of autism provides a great opportunity to shed light on the neural basis of atypical sensory experience and its possible interaction with social-cognitive domains in ASD.

Given that sensory hypersensitivity and enhanced detail-oriented perception are often reported in autism, we decided to investigate basic sensory processing and contextual modulation of different visual spatial frequency channels. Low SF channel may be responsive to the global patterns that can be used to learn regularities in the visual environment. On the other hand, high SF channel may be preferentially responsive to the visual details and could be important for thelocal processing of objects. Our behavioral results suggest that FX mice have reduced habitation and visual surprise adaptation compared to WT animals. Even though behavioral and neurophysiological data were acquired using different stimuli in oddball paradigms, we suggest that the observed behavioral deficits might arise from the discovered enhanced processing in late visual responses and reduced neural adaptation. Reduced habituation has been reported previously and linked to hyperexcitability and elevated stress in FX. However, reduced surprise habituation suggests that the extraction of common visual features might be altered in FX mice.

We demonstrated excessive visual processing in late neural responses in V1, especially at high spatial frequencies. This finding is consistent with the prior structural-functional imaging studies showing local hyperconnectivity within V1 and enhanced focus on details in the visual perception of autistic patients. It supports previous psychophysical and physiology studies showing altered spatiotemporal processing of high SF information in autism (Caplette et al., 2016; Kéïta et al., 2014). We also showed that units preferring SF distant from the target oddball SF of 0.03 cpd displayed reduced adaptation and enhanced PEs in FX vs WT animals. This observation may be explained by the reduced cross-frequency adaptation in FX mice. Alternatively, it might be an inherent feature of low and high SF channels. A direct comparison of STD and DEV responses between groups revealed differences mainly in the low SF channel. This finding suggests that mostly the low SF channel is affected by the reduced adaptation and enhanced prediction errors. Altogether, these sensory alterations would explain reduced habituation to common patterns/regularities and hypersensitivity to sensory stimuli.

One of the limitations of the SF oddball paradigm is that both STD and DEV stimuli have the same spatial frequency. PEs reported in this study were close to 0 or even negative for neurons preferring SF of 0.03 cpd due to the strongest adaptation in those neurons. It is important to note that all reported PEs would be an underestimation of true PEs due to the cross-feature adaptation. To address this issue, we investigated true PEs in units not preferring SF of 0.03 cpd. We found a reduced adaptation and enhanced prediction errors in FX mice across the cortical column. Furthermore, the SF population tuning curves of STD and DEV were altered in superficial and deep layers but not in layer 4. Higher firing rates to the SF tuning sequence were only found in the superficial layers of FX mice, which is in line with impaired sensory processing in superficial but not in layer 4 of somatosensory cortex in FX mice (Juczewski et al., 2016). Together, our findings suggest that laminar processing might be altered in V1 of FX mice.

The major rationale behind using spatial frequencies rather than more conventional orientated gratings is that it allows us to investigate the processing in both low and high SF channels, which effectively represent global and local visual features (Boeschoten et al., 2005; Hughes et al., 1996; Shulman et al., 1986). Combining data from tuning and oddball experiments allowed us to study both basic visual processing and contextual modulation of common and unique features. We find it striking that FX animals have excessive processing of high SF information and reduced adaptation in low SF channel suggesting that low and high SF streams might be differentially altered in FX mice. We think that our findings are aligned with reported sensory alterations in FX and autism (Deruelle et al., 2004; Rais et al., 2018).

In conclusion, our study provides evidence for altered processing in both low and high spatial frequency channels in FX. We extend prior research on visual perception in autism by mapping observed phenotypes to the specific neural computations that might underlie them. Early sensory processing is critical to building an internal representation of the external world during development. It also plays an important role during adulthood as animals continue to refine their internal model of the world o better fit the environment. It would be of great interest to study how alterations in different spatial frequency streams would affect higher-level cognitive processing in FX and autism.

## Author Contributions

A.P. and A.A.C. designed the study, A.P. and S.T.K. performed the experiments, A.P analyzed the data, A.P., S.T.K. and A.A.C. wrote the manuscript.

## Acknowledgments

We thank Sotiris Masmanidis for providing silicon probes, Maria Dadarlat for feedback on the manuscript, and Chubykin lab members for useful discussions. This work was funded by the National Institute of Mental Health (R01 MH116500) to A.A.C.

## Materials and Methods

### Experimental animals

All animal experiments were approved by the Purdue University Animal Care and Use Committee. The following strains were used to generate mice for this study: B6.129P2-Fmr1tm1Cgr/J (Fmr1 KO, JAX Stock No. 003025) and wild type (WT) C57/BL6. We used 10 male Fmr1 KO and 7 littermate controls. In total, we used 14 Fmr1 KO and 17 control animals for electrophysiology experiments. For pupillometry, we used 13 mutant and 13 WT mice after electrophysiological recordings. Animals were group-housed on a 12 hr light/dark cycle with full water and food access.

### Surgical procedures

Animal surgeries were performed as previously described (Pak et al., 2020). Briefly, about 2-month-old animals were induced with 5% isoflurane and secured to a motorized stereotaxic apparatus (Neurostar). Their body temperature was controlled using a heating pad and they were maintained at 1.5-2% isoflurane anesthesia. The scull was exposed to install a small head post and a reference pin. The binocular V1 coordinates (from lambda AP ±0.8 mm, LM: ±3.2 mm) were labeled using a Neurostar software with an integrated mouse brain atlas. Medical grade Metabond™ was then used to seal all exposed areas and form a head cap. After surgery, all animals were monitored for 3 days for any signs of distress or infection. Mice were then habituated to a head-fixation apparatus for at least 4 days 90 min per day. They were positioned in front of the monitor that displayed a grey screen. On the recording day, a small craniotomy was made above V1 on one of the hemispheres under 1.5% isoflurane anesthesia. They were then moved to the recording room and head-fixed to the apparatus in front of the monitor screen.

### Electrophysiology

All recordings were performed in awake head-fixed mice. After mice were transferred to the recording room, 30 min was allowed for mice to recover from anesthesia. After that, we inserted 64-channel silicon probe (Shobe et al., 2015a) (channel separation: vertical 25 μm, horizontal 20 μm, 3 columns, 1.05 mm in length) to perform acute extracellular electrophysiology. Each mouse underwent a maximum of two recording sessions (one per hemisphere). We acquired data at 30 kHz using OpenEphys hardware and software. We used an Arduino board to synchronize recordings and visual stimulus presentations using TTL communication. Custom written Python scripts using PsychoPy and pyserial were used to present visual stimuli and send TTL signals. Trypsin (2.5%) was used to clean the probe after recording sessions.

### Histology

Animals were anesthetized with 100 mg/kg ketamine and 16 mg/kg xylazine solution. Mice were then perfused transcardially with a 1x PBS followed by a 4% paraformaldehyde. After decapitation, their brain was extracted and stored in PFA in the fridge. After 24 hours, the brain was sliced in 0.1 mm sections in PBS using a vibratome. Coronal slices were mounted on slides using n-propyl-gallate media and sealed with transparent nail polish. Slices were imaged using light microscopy (VWR) to verify the probe placement in V1.

### Visual stimulation

We used a PsychoPy, an open-source Python software, to create and present all visual stimulations (Peirce, 2009). A gamma calibrated monitor (22’ ViewSonic VX2252, 60 Hz) was used to present visual stimuli. The mean luminance of the monitor was 30 cd/m^2^. The monitor was placed 17 cm in front of the mouse to binocularly present stimuli. To generate visual stimulations for a spatial frequency tuning and an oddball paradigm, we performed a spatial frequency filtering of random noise. Specifically, we band-pass filtered random noise in different non-overlapping SF bands. This was done by performing the following steps. First, we randomly generated noise and converted it to a frequency domain using FFT (numpy FFT). Second, we created a spatial frequency bandpass filter using psychopy Butterworth filter with an order of 10. Third, we multiplied the white noise in the frequency domain by our band-pass filter. This step filtered all the frequencies but the desired SF band. Fourth, we took the inverse Fourier transform of our altered frequency domain. The procedure and a Python code for spatial frequency filtering were adapted from http://www.djmannion.net/psych_programming/vision/sf_filt/sf_filt.html. We modified the above code to generate SF filtered noise. Overall, we used 6 different spatial frequencies for SF tuning: 7.5E-3, 0.015, 0.03, 0.06, 0.12, and 0.24 cycles/degrees. We chose these frequencies based on previous studies and known spatial frequency tuning of mouse V1 neurons. We verified that we can obtain reliable SF tuning in this and our previous study (Kissinger et al., 2018). SF tuning sequence contained 6 different SF stimuli presented in a pseudorandom order at equal probability. We used an inter-trial interval of at least 4s to prevent any adaptation. Furthermore, SF filtered stimuli were randomly generated on each trial to uniformly sample different receptive fields. This was mainly important for lower spatial frequencies. For the oddball paradigm, we used two stimuli of the same SF but the different overall pattern. The first stimulus was a standard (STD) with a probability of 0.875. Its texture did not change across trials. The second one was a deviant (DEV) with a probability of 0.125, its overall pattern changed across trials. This was done to maximize the surprise response. Inter stimulus interval was randomly chosen from the range of 0.5 and 1.2 s. The stimulus was presented for 0.5s.

### LFP analysis

Raw electrophysiology traces were first downsampled to 1 kHz. We then used symmetric linear-phase FIR filter (default parameters) from the mne Python library to remove 60Hz noise. Next, we identified a Layer 4, by finding a channel with the strongest negative deflection in the first 100ms after stimulus onset. Time-frequency analysis was done using a complex wavelet convolution. 40 different wavelets were designed across a logarithmic range of 2-80 Hz, with cycles ranging from 3 to 10. This gave us an optimal time-frequency precision tradeoff. We convolved these wavelets with averaged LFP traces and then averaged the resulting power spectra across different conditions. For heatmaps, power was dB baseline normalized. To quantify a mean power within a particular band, we averaged responses within a 0.35-0.5s time window. We used 6 different frequency bands: theta (4-8Hz), alpha (8-12 Hz), beta (12-30 Hz), low gamma (30-50Hz), and high gamma (50-80Hz). CSD analysis was done using spline-iCSD method implemented in Python (Pettersen et al., 2006). We used default parameters but number of steps was changed to 41.

### Single unit analysis

Clustering and manual curation of units were performed as previously described (Pak et al., 2020). Kilosort was used for a spike detection and sorting. It uses a template matching algorithm and allows a GPU acceleration (Pachitariu et al., 2016). Default configuration parameters were used for clustering but a threshold for spike detection was changed from −4 to −6. SD. Templates were initialized from the data. Kilosort was run using MATLAB (Mathworks) on Windows 10 running machine with an NVIDIA GeForce GTX960 graphics card. For clustering purposes, all the different recording blocks were concatenated together. This allowed us to track single neurons across different recordings sessions. After clustering, we visualized and verified clustering results using Klusta/Phy GUI. It speeds up the process of manually removing, splitting, and merging units (Rossant et al., 2016). We used several criteria to only include well isolated units: 1) had more than 100 spikes for each experimental block, 2) less than 5% of spikes violated an absolute refractory period, 3) clean template shape, and 4) templates are localized within a small channel group. To merge and split units, we followed the guidelines available online (https://github.com/kwikteam/phy-contrib/blob/master/docs/template-gui.md). Peristimulus time histograms (PSTHs) of single units were constructed by binning spike times across trials with 10 ms bins and convolving the obtained histogram with a Gaussian Kernel (width = 100 ms). Z-score was calculated by the following formula:

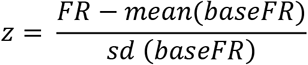

where FR is a firing rate at each time point and base refers to the baseline activity over 0-0.3s. We also computed modulation indices for SSA and PEs.

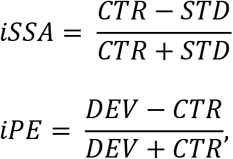

In this study, we focused on neurons that upregulate their firing in response to visual stimuli. We used Wilcoxon Signed rank test to identify these neurons by comparing baseline firing rate 0.05-0.35 versus stimulus window 0.35-0.65s. For spatial frequency tuning analysis, we first found preferred SF for each neuron. It was defined as the maximum of baseline (0-0.3) subtracted firing rate (0.35-0.5) in response to different SF bands. Population tuning curve was then constructed by either averaging firing rate or first rescaling and then averaging across neurons. In the latter case, each neuron gets an equal weight. The response to the SF0.03 was used as the control for oddball paradigm. To equalize the number trials between STD and DEV stimuli, we only used pre-DEV trials for STD. These units were then further split by the cortical depth. Layer of each neuron was assigned based on the depth of the channel with strongest negative deflection of the template.

SF neural decoding was performed using Linear Discriminant Analysis in Python scikit-learn package (default parameters). Population spike counts from different time windows were used to train classifiers. We used a 4-fold cross-validation with 5 repeats. The number of folds was chosen so that the test size was not below 30 samples. We also tried to train logistic regression (multinomial) and SVM (with RBF kernel) classifiers but LDA gave better performance given the number of parameters to specify.

### Pupillometry

Video acquisition and analysis was performed as previously descried (Kissinger et al., 2018). Briefly, we acquired mouse pupil videos under IR illumination. Videos were then analyzed post hoc using a Python computer vision library, OpenCV. One of the key components of the preprocessing was a histogram equalization, which improves the contrast of the videos. We then thresholded frames to only include pixel values below 15. Given a good preprocessing pipeline, the pupil tracking was performed by first detecting contours and then fitting a minimum enclosing circle. This ensured that whiskers and small local contrast variations did not affect the tracking. We then extracted x,y-coordinates, and radius of the circle. We analyzed both a raw diameter of the pupil and area % change from the baseline. Raw pupil size was used to study habituation across oddball blocks. Baseline normalized responses were used to compare surprise response between different sessions.

### Statistical Analysis

We used scipy.stats Python library to perform statistical analysis. Data were not tested for normality of residuals and only non-parametric tests were used. Mann-Whitney U test was used to compare two independent populations. It was used to compare a trial-averaged LFP and neuronal firing rate in response to different SF bands WT vs FX; responses to STD, DEV, and CTR in both LFP and units. A total nine comparisons were tested. 3 within each group: STD vs DEV, STD vs CTR, and DEV vs CTR (WT and FX) and 3 between groups within each stimulus condition. P-values were adjusted using a Benjamini-Hochberg procedure that controls for a false discovery rate. Kolmogorov-Smirnov 2 sample test was used to compare distributions of iSSA and iPE indices between WT and FX mice in different layers. Paired t test was used to compare pupil baseline diameter and surprise response between session 1 and 2 in WT and FX mice.

### Data and code availability

The data that support the results of the current study and the analysis code are available from the corresponding author upon reasonable request.

